# Progressive Sperm Separation Using Parallelized, High-Throughput, Microchamber-based Microfluidics

**DOI:** 10.1101/2020.07.31.231373

**Authors:** Mohammad Yaghoobi, Morteza Azizi, Amir Mokhtare, Alireza Abbaspourrad

**Author notes:** **Author Contributions**, M.Y. and M.A. designed the platform; M.Y., M.A. and A.M. performed the research; M.Y. did the simulation and mathematical models, M.Y., M.A. and A.A. analyzed the data; M.Y., M.A., A.M. and A.A. wrote the paper; A.A. was the principal investigator of the group.

## Abstract

Motility is one of the most important factors in sperm migration toward egg. Therefore, sperm separation based on motility increases the chance of the best sperm selection in the process of infertility treatments. Unfortunately, it is now vastly done by conventional procedures which lack certain delicacy and precision and increase the risk of damage to sperm cells. Microfluidic systems, on the other hand, can sort sperm in a less intrusive way. However, microfluidic techniques have yet to receive widespread adoption in clinical settings, not only due to their relatively cumbersome operation, but also their extremely low outcome, leaving them inefficient in practice. Here we propose a microchamber-based microfluidic platform that can separate progressive motile sperm from nonviable sperm and debris as well as trapped nonprogressive sperm in the microchambers. Our platform is operated in a short period of time (<10 min) with an excellent degree of controllability, without any prior sample preparation. Our results show that the microchambers’ depth does not affect the residence time of motile sperm. Therefore, we are able to inspect high sample volumes (1 mL) within the same time. Furthermore, we maximize the concentration of the collected sperm by tuning the washing medium flow rate above the sperm rheotactic threshold. We foresee that our microfluidic platform may provide a facile solution for high-throughput, robust, and easy-to-modify for collection of progressive sperm needed for assisted reproductive technologies (ARTs).

**Significance Statement:** Assisted Reproductive Technologies require efficient, minimally invasive, and fast methods of sperm separation. Centrifugation methods used in clinics and biological research labs, fall short in these aspects as they are low-yield, intrusive to sperm’s DNA, and time consuming. We have developed a microchamber-based microfluidic platform for high-throughput separation of progressive motile sperm from undiluted raw semen samples. The method was further optimized to increase the concentration of collected samples. Higher concentration of collected samples combined with higher motility of the separated sperm compared to those in raw semen, make it a suitable choice in clinical applications, fertility diagnostics, and fundamental research.

## Introduction

Sperm quality is central to the success of Assisted Reproductive Technology (ART), as it can directly alter the fertilization outcome and offspring health (1, 2). Sperm separation plays a crucial role in improving sperm quality, both in research and clinical settings, needed to discover the underlying factors behind infertility. DNA integrity, hyaluronic acid binding capability, motility, morphology (2) capacitation ability (3), and plasma membrane negative charge (4) are different sperm characteristics that have been evaluated to this date. Among these, progressive sperm motility is a critical factor that enables sperm to pass through barriers in the female reproductive tract (5, 6).

Conventional methods of sperm separation, which mostly focus on sperm motility, are gradient based centrifugation and swim up. (1, 7). However, these methods have detrimental effects on the sperm’s genetic load (8, 9) and cause an increased rate of apoptosis in infertile men’s sperm (10). This shows both the flaws of these methods, and the sensitivity of the infertile couple’s sperm compared to fertile males. In addition, these methods require a stringent skill set, lack standardized procedures, and prone to human errors (1), not to mention that they unfavorably are time consuming. However, new technologies with minimal perturbation, such as microfluidics, have been extensively explored and considered as promising remedies for the flaws associated with the conventional methods (2, 7, 11–15).

Among all microfluidic platforms used for sperm separation, rheotaxis (16), crossing the streamlines (14), random motion (11), and boundary-following inclination (15, 17) of the motile spermatozoa are the most popular ones. Rheotaxis, which is a passive, hydrodynamically controlled process (18), was used in a diffuser type microfluidics device to select motile sperm (19). Apparent random motion of sperm in a quiescent medium (which results in diffusion) has been guided within space-constrained microfluidic channels to separate racing sperms (20). Persistence length of collected sperm was increased in a similar design by addition of pillar arrays (12). The propensity of progressive motile spermatozoa to move along surfaces and near the corners was used to design a microfluidic platform with a loading ring and parallel walls to guide human motile sperm to the outlet (1, 15, 21).

Sperm separation time is an important factor since it can compromise the sperm motility, viability, and DNA integrity if it takes more than 2 hours (22). Reducing the time of sperm separation is proved to be effective in the of Intracytoplasmic sperm injection (ICSI) (23). One estimation of processing time is the interrogation of 1 mL sperm sample in 10 minutes (1).To the best of our knowledge, none of the previous methods (7, 11–15, 20, 24), which used sperm motility feature for sperm separation, meet this criterion. Here, we propose a microchamber-based microfluidics platform for sperm separation using boundary-following ability of motile sperm with the least possible invasiveness as well as dramatic increase of the interrogation rate of raw semen samples. The array of microchambers can be readily parallelized into a large network (*e.g*., a few thousand microchambers) without changing the processing time. Therefore, we can increase the volume of interrogated samples (up to several milliliters of raw semen sample) within the same 10-minute processing time. More importantly, the initial sample does not need any dilution or preprocessing (as opposed to the previous methods). It thus enables us to interrogate raw semen samples right after ejaculation and collection from patients.

## Results and Discussion

### Microfluidic Device and Procedure

The microfluidic device consists of a micropatterned polydimethylsiloxane (PDMS) layer, fabricated using soft-lithography technique and plasma bonded to a glass slide. The PDMS layer entails parallel microchambers that are perpendicularly connected to longitudinal main channels with connecting channels. The device has two ports for sample loading and discharging. The schematic arrangement of the main channels and microchambers, along with scanning electron microscopy (SEM) and photographs of the device are shown in Fig. 1A and 1B.

**Fig. 1.**
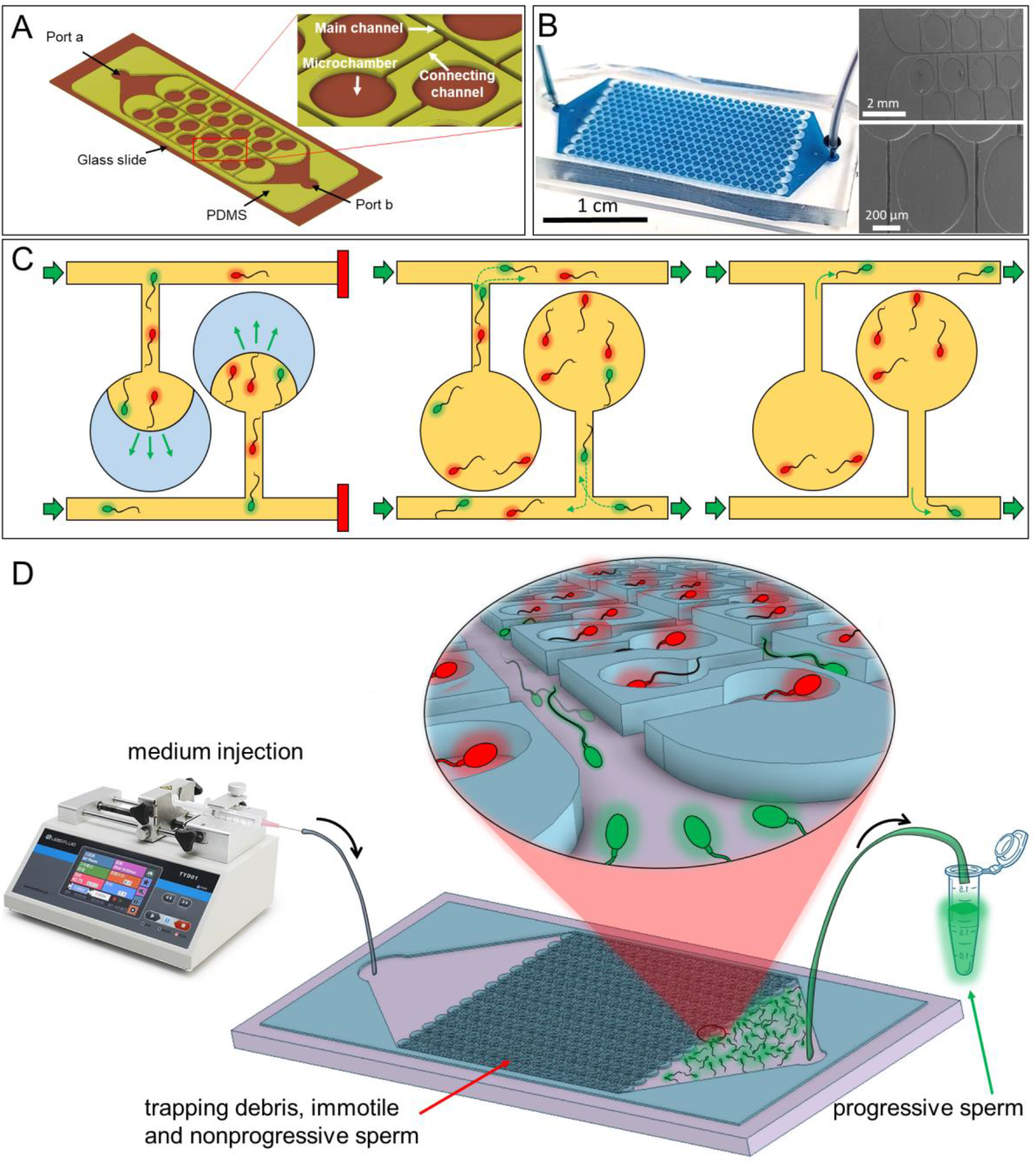
(A) Schematic representation of the platform design for sperm separation. (B) The photograph of microchambers filled with blue dye. (C) Loading of the semen sample into the microchambers, air discharge because of porosity of PDMS and washing the main channel for sperm collection. (D) The parallelization capacity of the platform for increasing the volume of loaded sample and simplicity of the collection process.

Device is operated in three steps: First, it is loaded with a raw sample. To do so, we the outlet (port -b) is temporarily blocked while the sample is pushed into the device from the inlet (port a). Due to PDMS porosity, trapped air inside the microchambers is discharged, allowing the raw sample to gradually fill them. Second, its main channels are washed with a fresh medium. In this stage, the outlet is reopened and fresh media is injected into the main channels, clearing them from any debris and non-viable sperm. Third, the fresh medium is gently injected from the inlet into the main channels and progressive motile sperm are collected at the outlet. During the washing step, motile sperm naturally navigate along the microchamber walls (boundaries) that automatically guide them toward the main channels. Upon reaching the main and connecting channels intersections, motile sperm are washed away toward the outlet and collected. The collecting step is continued until the majority of the motile sperm find their way out of the microchambers to the main channels. Device operation steps are illustrated in Fig. 1C.

To optimize our device, we investigated the impact of channel depth and microchamber diameters on the loading capacity and separation performance of the device. In doing so, we changed the depth of the microchannels from 30 μm to 70 μm. In addition, we varied the microchambers diameters (350 μm, 500 μm, and 1000 μm) to evaluate the effect of boundary length on the separation performance of the device. Detailed illustration of device dimensions can be found in Fig. S1 and Table S1.

It should be noted that the overall processing time remains intact even for increased volumes of samples resulted from increasing the number of microchannels and changing the channel depth. This is mainly because the collection time is directly a function of microchamber diameter and the time that a motile sperm takes to travel along its boundary to exit (called residence time). The time for washing step is only a fraction of sperm residence time in the microchambers and does not significantly affect the total separation time.

Overall, the device operation is simple and only required a syringe pump, a syringe for injecting fresh medium, and an Eppendorf tube to collect the progressive motile sperm from the outlet, as illustrated in Fig. 1D. Thus, this cheap (cost ~ $2/device), easy-to-operate, and high-throughput platform can be an ideal and non-invasive alternative for separation of progressive motile sperm needed for clinical and research studies.

### Device Loading Time and Volume

As the chambers’ diameter increases, the device loading time increases accordingly. Theoretically, microchambers’ loading time with a raw semen sample is a function of PDMS porosity, applied pressure (25) (or mass flow at the inlet), and microchamber volume. In *SI section 2*, we used simplifying approximation and showed that loading process can be theoretically modeled by a set of equations. In Fig. 2, sample progression inside the microchambers is visualized through images taken from a time-lapse. Dashed red lines are the common interface of the sample and air trapped in the chambers. The effect of three different microchamber diameters (350 μm, 500 μm, and 1000 μm) and two different flow rates (540 μL/h and 1240 μL/h) on the loading time is shown in Fig. S3. Similar to our theoretical predictions, experiments showed an exponential correlation over time for loaded volume. In other words, the rate of loading is initially fast but decreases as the loaded volume inside the microchambers increases until the microchamber is fully loaded. More importantly, loading occurs simultaneously and almost uniformly for all the microchambers (as indicated by the narrow error bars in Fig. S3). This feature gives us the ability to increase the number of microchambers to millions while having a robust control over loading times.

**Fig. 2.**
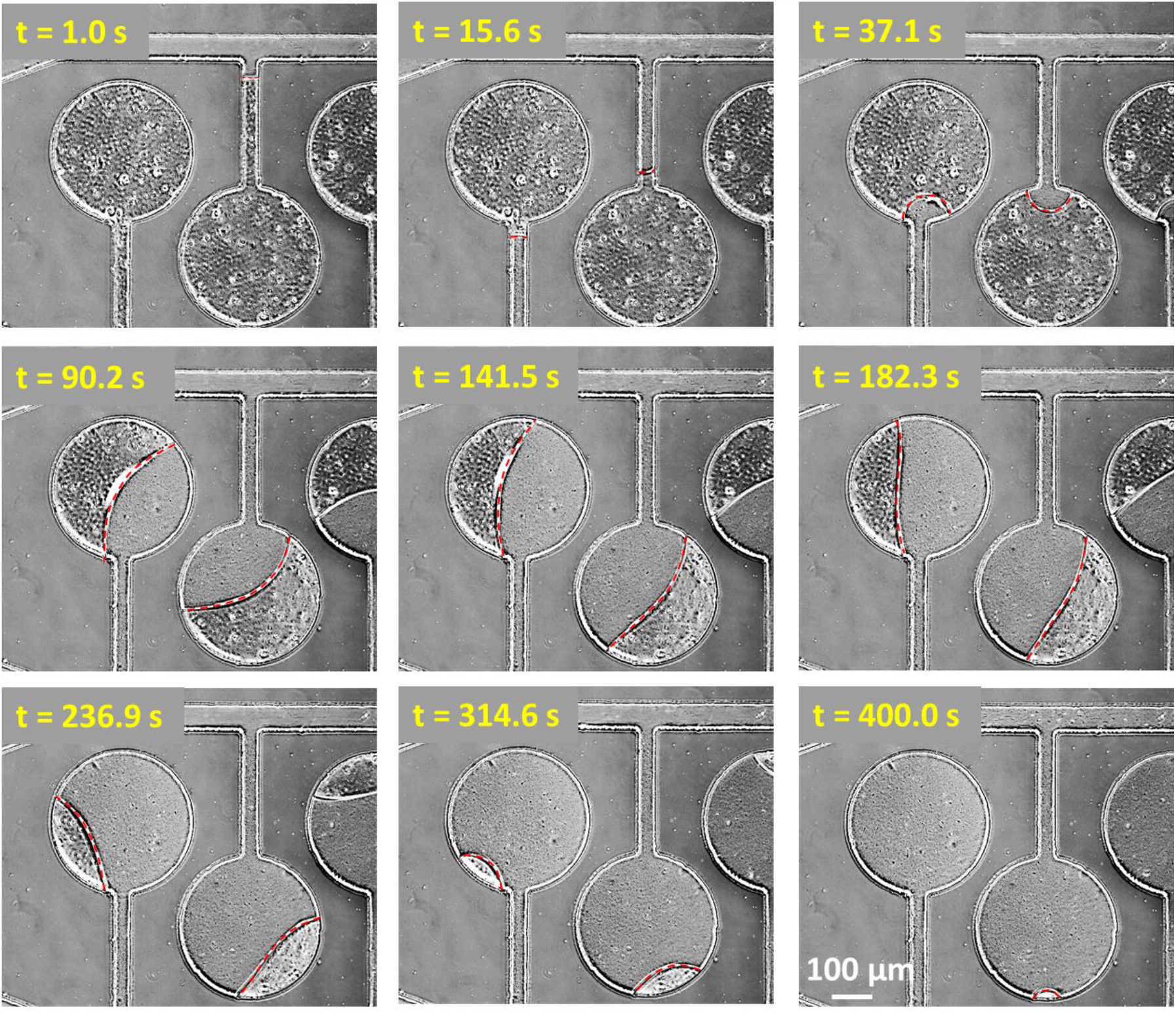
The loading process for microchambers with “350 (D) × 70 (H) μm^2^” dimensions. The common interface of the sperm sample and the air trapped in the channel (dashed red line) is attenuated in each time-lapse. This experiment was performed by applying constant pressure on the sample loaded into the chip while the outlet of the main channel is closed.

### Sperm Number Equilibrium in the Microchambers

With successful loading of the raw semen sample into the device, progressive motile sperm show everywhere in the microchip including the microchambers and main channels. Due to their motility, they constantly move from the main channel to the microchambers and vice versa. Such movements quickly settles into an equilibrium between the sperm that move into microchambers and those who leave it, resulting in an average motile sperm population inside the microchannels. Reaching this equilibrium is crucial as it later provides us with a high degree of flexibility in washing the main channel and removing all the debris (including motile and immotile sperm). We validated our hypothesis by observing an intersection of a connecting channel with the main channel as demonstrated in Fig. 3A. As it can be seen in the time-lapse images, two spermatozoa enter and exit the microchamber (shown in red and green). The difference between the cumulative population of sperm exiting a microchamber and those entering it remains constant over 400 seconds, indicating that an equilibrium state remains dominant over a long period of time, (Fig. 3B). More importantly, the rate of sperm exchange between the microchambers and the main channels is 0.08 1/sec, which is small enough to provide an operator with ample time to prepare for the washing and collection steps. The details of sperm trajectories in the vicinity of an intersection are demonstrated in Fig. 3C, where purple dots represent sperm exiting the microchamber and red and blue dots represent sperm swimming toward the microchamber entrance from the right and left walls. Not all the sperm approaching the intersection enter the microchamber that is highlighted by the intensity of the blue and red colors in the left part of Fig. 3C. Darker colors represent sperm entering the microchamber and lighter colors represent sperm that are deflected from the connecting channel and do not enter it. Our results showed that nearly half of the sperm swimming from left and right of the connecting channel entered the chambers (dark blue and red dots in Fig. 3C). The main reason behind this phenomenon is that sperm who approach the intersection with the right angle will detach from the main channel wall at different deflection angles, resulting in swimming directions toward or away from the microchamber (26, 27). Our measurements show that the deflection angle for the right and left swimmers are 22.19° (± 13.5°) and 21.51° (± 13.2°), respectively, which are consistent with the reported scattering data for the bull spermatozoa at 29 °C (27). Results also show that sperm deflection angle is independent from the swimming direction, and that sperm populations entering from the right and left walls are almost equal. Also, sperm exiting the microchambers did not show any preference for swimming direction. In other words, sperm exiting the microchambers toward the front wall have a 50% chance of swimming to the left or to the right. Furthermore, the number of the sperm approaching the intersection from right or left is approximately the same. In short, our results show that this microchamber-based platform reaches a kinematic equilibrium state, in terms of sperm count, that is highly preserved.

**Fig. 3.**
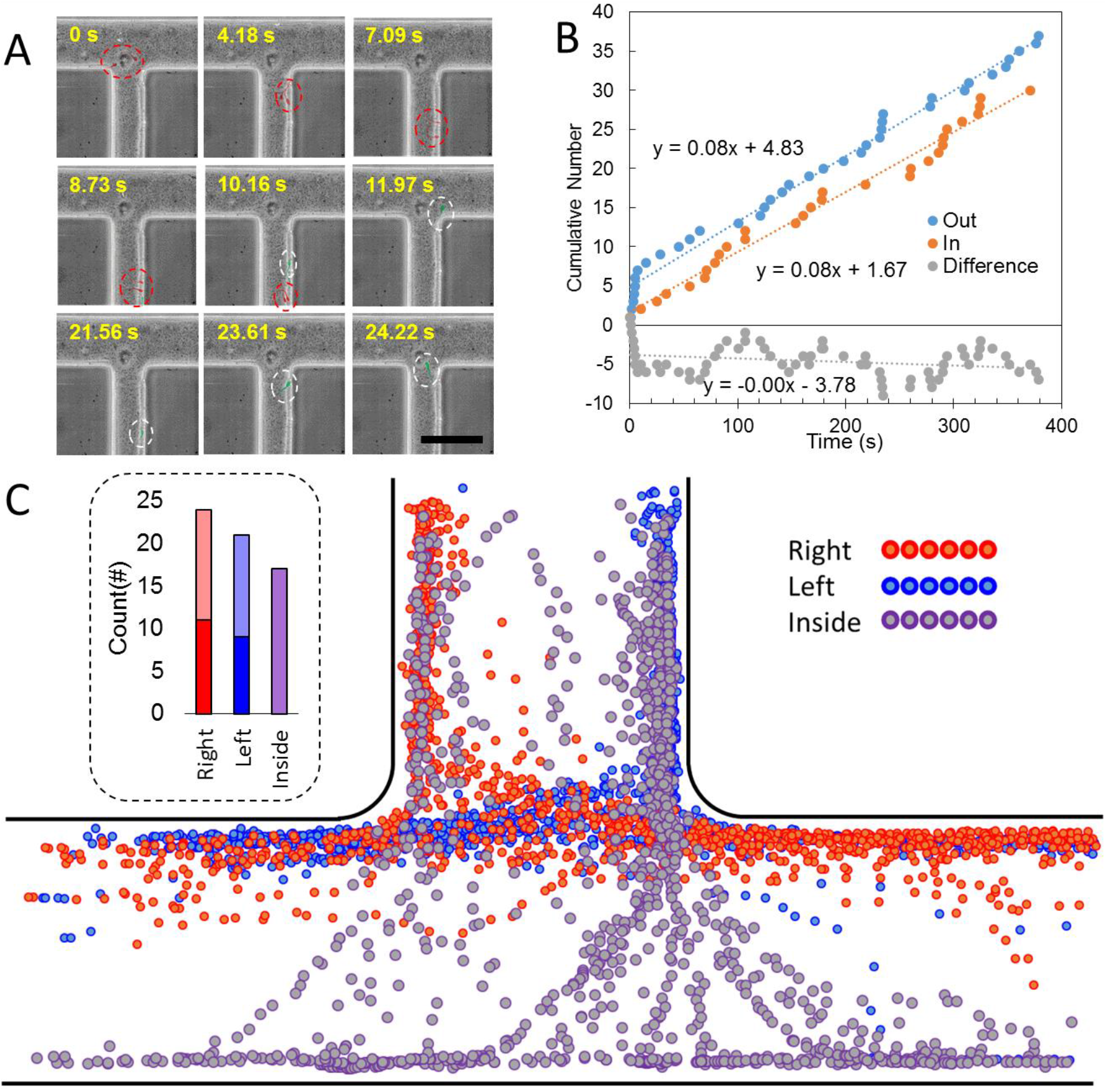
The overall number of the sperm in the chambers remain constant. (A) Different time lapses of a typical junction at the equilibrium state shows two sperm going into the channel (highlighted as red color and put in red dashed curves) and two other sperm coming out (highlighted as green and encircled with white color curves) in the course of 24 seconds. Scale bar is 100 μm (B) Cumulative number of sperm going in and out of a chamber in a 400 second period. The slope of the fitted line for those outward and inward sperm is equal, bringing an equilibrium to the number of the sperm in the chamber. (C) Dot trajectories of the sperm swimming near the junction wall from right (red dots), left (blue dots) and those coming out of the chamber swimming near the connecting channel walls (purple dots). Number of these sperm is shown in the left diagram with the same color code. The light blue and red colors show the sperm that go into the chamber from right and left, respectively.

### Extraction and Residence Time in Collection Step

Motile spermatozoa trapped inside the microchambers during the loading step actively interact with their surrounding boundary and swim toward it. Based on their incident angle with the microchamber wall, they either navigate in a clockwise or counterclockwise manner along the boundaries (28) until they are guided toward the main channel. Upon entering the main channel, they are washed away with a flow of the fresh medium, which is continuously injected, and collected at the outlet into a small tube. Hydrodynamics dictates that fresh medium only flows through the main channels where even higher injection speed doesn’t disturb the quiescent conditions (i.e., dead zones) inside the microchambers (Fig. 4A.). This is important as it not only provides the right conditions for motile sperm navigation toward the washing flow but ensures the entrapment of immotile particles/sperm inside the microchambers. This is illustrated both experimentally and numerically using an injection flow rate of 10 μL/h from the main channel’s inlet (left and right side inset of Fig. 4A, respectively).

**Fig. 4.**
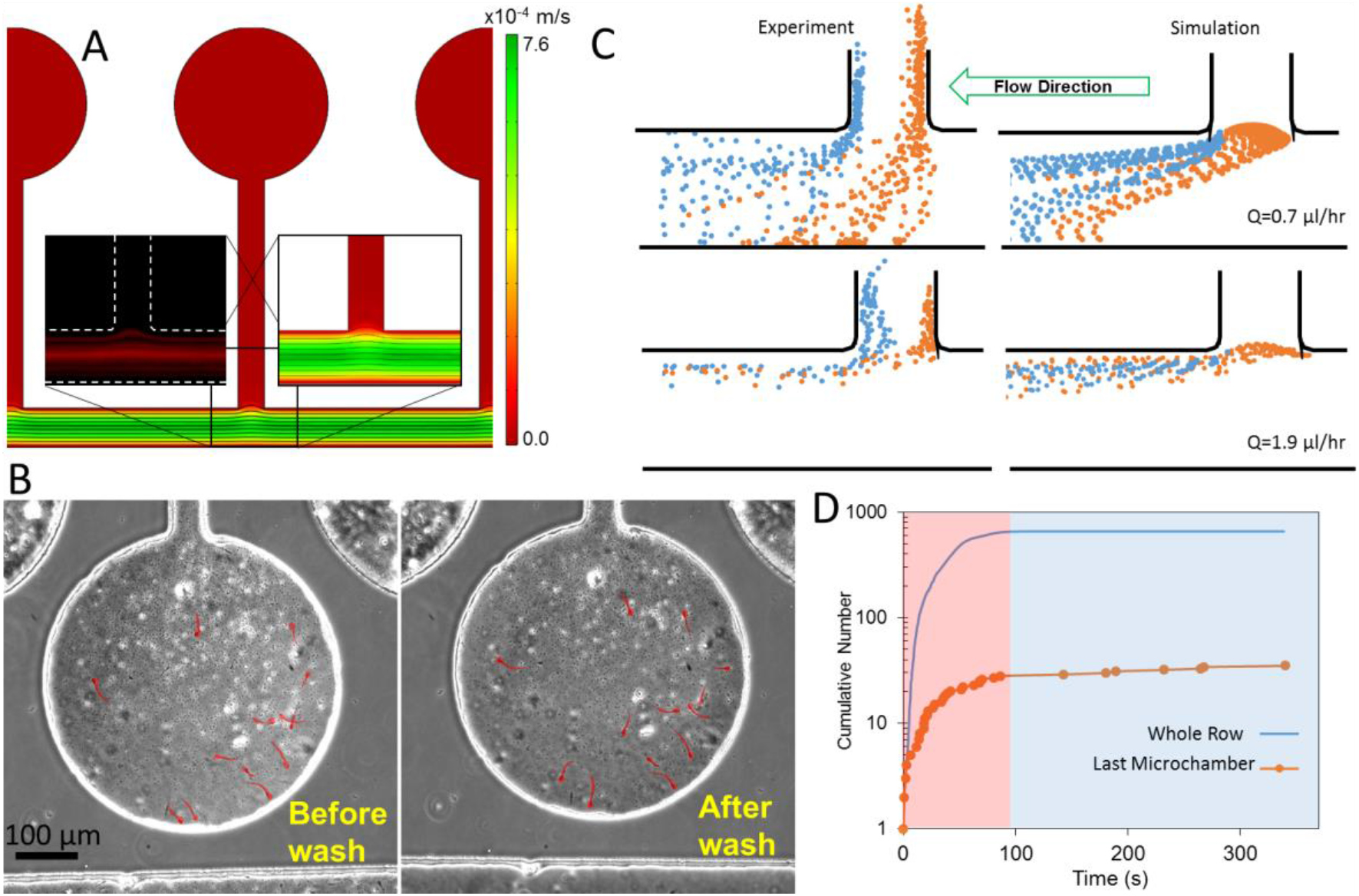
Washing steps and sperm extraction. (A) the streamlines and the velocity magnitude in the main channel for volume flow rate of 10 μL/hr. The streamlines of the magnified region is shown experimentally with fluorescent particles. (B) the dead sperm and particles remain in the microchamber. The dead sperm are highlighted in red. Scale bare is 100 μm (C) Rheotaxis for both simulation and experiments (D) typical extraction diagram of the device over time for 500×30 μm^2^ of the last chamber in a row and whole number of sperm in that row.

Fig. 4B shows the position of the immotile sperm loaded into a particular microchamber before and after washing and collection steps. As can be seen, the positions of dead sperm remain unchanged or only slightly changed due to their interaction with motile sperm. It is noteworthy to point to the absence of motile sperm in the “before washing step” image (Fig. 4B, left), because motile sperm are only present alongside the walls of the microchamber that are under the halo-scattered light effect of microscopy.

Regarding the platform efficiency, the concentration of motile sperm collected from the outlet is inversely proportional to the injection flow rate of the fresh medium. Higher injection rates do not improve the performance and only drop the final concentration of motile sperm due to excessive amount of medium used during the washing step. The collected sperm volume should not exceed the initial raw sample volume (loaded to the device), otherwise, collected samples would be too diluted and require centrifugation which defies the purpose of the microfluidics-based separation. On the other hand, lower injection flow rates—below an optimum value—increase the chances of sperm rheotaxis (a behavior that is not desired) inside the main channels. With Injection flow rates lower than the sperm rheotaxis velocity, progressive sperm can swim in the opposite direction of the injected medium and enter to the adjacent microchambers, thus increasing the collection time as well as lowering the microchip efficiency. Note that rheotaxis only occurs in a narrow range of mass flow rates (7, 29) and flow rates greater than this range, overcome sperm’s swimming force and easily drag them downstream. In this regard, we investigated the effect of flow rates on the rheotaxis of sperm exiting the microchambers experimentally and numerically. Fig. 4C shows the sperm trajectories are distorted by the medium flow at different flow rates (Q = 1.9 μL/h and 0.7 μL/h.) compared to Fig. 3C when there is no flow. Dot trajectories of sperm swimming along the left wall of the connecting channel are shown in orange and those that are navigating near the right wall are shown in blue. Higher flow rate (i.e., Q = 1.9 μL/h) outweighs sperm’s motility and easily drags them toward the outlet, resulting in a 100% recovery of sperm swimming out of microchambers after the start of the collection step. Conversely, sperm’s motility becomes significant in the lower flow rate (i.e., Q = 0.7 μL/h), such that sperm can swim upstream swapping between the main channel walls in a butterfly shaped trajectory as reported previously [14]. Sperm swimming in such a pattern will find their way back to microchambers.

For numerical simulations shown in Fig. 4C, we used a population of 40 sperm with normally distributed velocities. For initial conditions, we released half of the sperm from an arbitrary point near the right-hand wall and the other half from an arbitrary point near the left-hand wall in a perpendicular direction to the flow. We used turning dynamics in sperm swimming direction (30) to calculate their 2D trajectories while they exit the chambers and are being exposed to the flow in the main channel. (The details of these simulations can be found in *SI section 4*).

Motile sperm collection step starts with pure medium injection into the main channels until all available progressive spermatozoa exit the microchambers. Fig. 4D compares cumulative sperm collection from the last microchamber (size 500×30 μm^2^) in a row of 48 microchambers to the total cumulative number of spermatozoa, collected from that specific row. As is evident, 28 out of 36 sperm (~ 78%) are collected within 100 seconds into the washing step and the remaining sperm need more than 200 seconds to exit the microchamber. A total of 655 sperm were collected in 300 seconds and the microchambers were checked afterwards to ensure no progressive swimming sperm remained in the microchambers. The samples are inhomogeneous in terms of the number of sperm in the microchambers, meaning that several trial results in various numbers of sperm extracted from a particular microchamber (*SI section 5*).

The residence time of loaded progressive sperm varies based on their initial position, velocity, swimming direction, the microchamber diameter, and the length of the connecting channels. For sperm with the same velocity, the minimum and maximum residence times belong to the progressive sperm which move along the shortest and longest paths from their original positions inside a microchamber until they reach the intersections that connects the microchambers to the main channels.

Accordingly, we neglected sperm-sperm interactions, assuming the movement of one sperm does not affect the motion path of its nearby sperm. We derived a formula for residence time distributions of a population of the loaded progressive sperm with respect to their velocity distribution. First, we considered a pool of progressive sperm with Gaussian velocity distribution <*v*, *σ_v_*> loaded into a microchamber with a known diameter (D) and a connecting channel with a known length (L). The distribution of the residence time of this population can be described by Eq. (1):

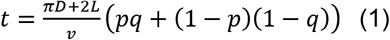

where p is the uniformity of the initial sperm distribution on the microchambers’ wall and q is another uniformly random variable set to be 1 for clockwise swimming sperm (half of the population) and 0 for the other half (counterclockwise swimming sperm). It should be noted that our calculations were done assuming that the system is at equilibrium where most of the motile spermatozoa have already reached the microchamber boundaries (more discussion on the topic can be found in *SI section 5*). Notice that this equation does not consider sperm collisions on the boundaries and assumes continuous sperm navigation along the boundaries upon their arrival at any contact angle. The residence times for all of the designs were calculated for a sperm population with v=80 μm/s and *σ_v_*=20 μm/s.

Theoretical and experimental data are plotted and shown in Fig. 5. The measured residence times resemble a left-skewed distribution and are in accordance with the theoretically predicted values. In addition, residence time change with increase in microchamber diameter is more notable for D = 500 to D = 1000 μm than D = 350 to 500 μm. Deviation of the predicted results (based on the model) from the experimental ones are also more significant for larger microchambers. We think this is mainly due to the fact that as the microchamber diameter increases, so do the sperm-sperm interactions which we assumed insignificant in our model. It is also clear that the residence times are independent of the microchip depth. This is because of the sperm accumulation on the top and bottom surfaces of the microchip and their preference for 2D navigation near top and bottom surfaces (31). The collection time for more than 90% of the sperm in the microchambers were 60, 135, and 215 seconds for D = 350, 500 and 1000 μm, respectively, and did not substantially vary when we increased the microchip depth from 30 μm to 70 μm.

**Fig. 5.**
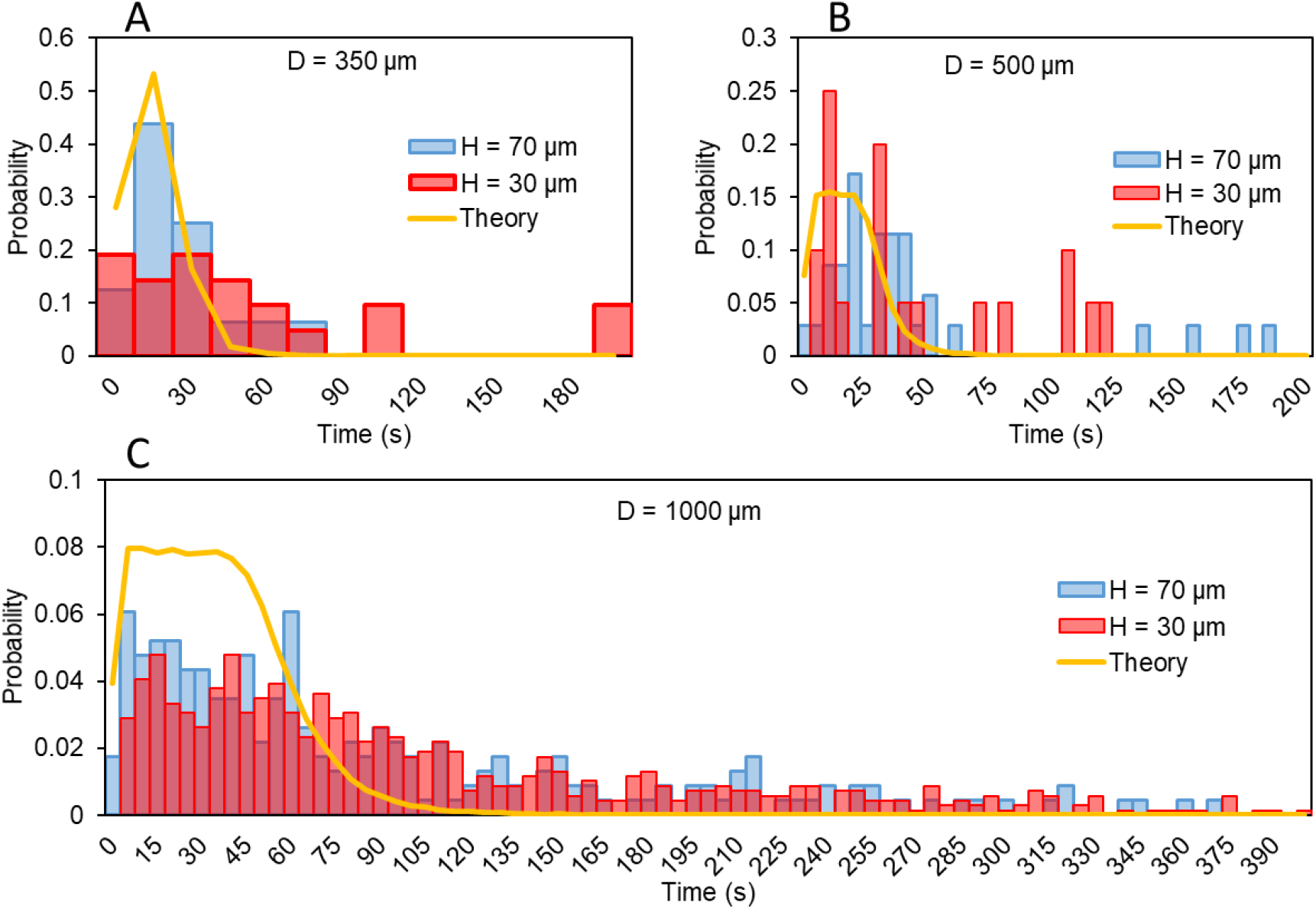
The distribution of residence times of sperm in microchambers with different diameters and 30 μm and 70 μm depths (the theoretical distribution is also shown for comparison).

### Characteristics of the Collected Sperm

The collection zone lies right before the outlet of the microchip (Fig. 6A). Due to the channel width expansion in this area, medium flow velocity significantly decreases, causing progressive sperm to undergo rheotaxis, although, they are not strong enough to return to the main channels. After the collection is done, the inlet flow is increased so that all the sperm are pushed to the outlet and collected in the collection tube. We further analyzed the progressive sperm accumulated in the collection zone by measuring and comparing their straight-line velocity (VSL) and average-path velocity (VAP), as shown in Fig. 6B. Results show a significant increase in the mean VSL from 60 μm/s in raw semen sample to ~80 μm/s in the extract due to the higher concentration of highly progressive motile sperm in the separated sample.

**Fig. 6.**
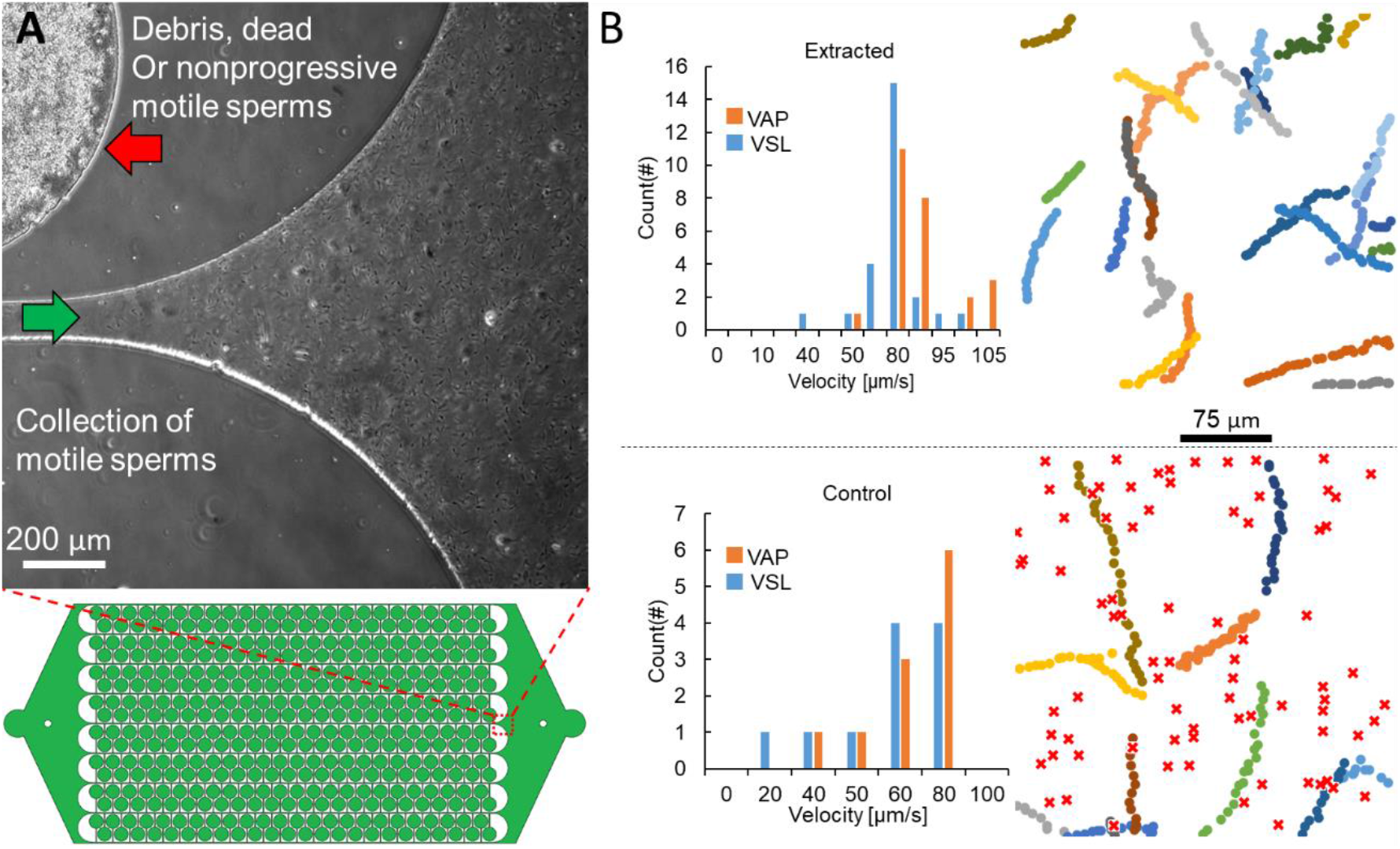
Bulk separation of motile sperm within the 1000×70 μm^2^. (A) The population of the sperm at the outlet is visible. Green flash shows the direction of washing sperm to the outlet and the debris, dead and nonprogressive motile sperm remain in the chambers. (B) The extracted population of sperm from the device has higher motility and concentration and far fewer number of dead sperm. VAP and VSL distribution of the extracted sperm and the control sample are illustrated. The trajectories of motile sperm are shown in different colors and dead sperm are shown in red crosses. (Debris which is present in the frozen thawed sample of bovine semen is also filtered out through the device and cannot be seen in the separated sample.

Furthermore, our proposed microfluidic platform effectively differentiates and separates the progressive and nonprogressive motile sperm. Our observations of a microchamber after 15 minutes revealed the formation of moon-like crater-shaped hollows as a result of circular motion of nonprogressive motile sperm. The circular movement of nonprogressive motile sperm pushes the debris (egg yolk extender particles) toward the microchamber walls and randomly creates hollow patterns inside the microchamber (Fig. S7). This fact combined with the absence of any circular motion within the collected sample are indicative of the capability of our platform to separate progressive motile sperm from the initial raw sample and trap nonprogressive sperm in the microchambers.

We also studied the effect of the size of the microchamber on the retrieving efficiency of progressive motile sperm from the microchip, as shown in Fig. S8A. To test the efficiency of these devices, we used raw semen samples with the concentration ~ 8.2 M/mL, of which 21% of the sperm were motile. The collected extracts from the 1000×30 μm^2^ and 500×30 μm^2^ devices showed approximately 80% retrieval of the progressive motile sperm from the initial sample. The retrieval efficiency was only 15% for the smallest device with the smallest diameter of testing microchambers (350×30 μm^2^). The low retrieval efficiency for 350×30 μm^2^ device is because of the short residence time of the sperm in the microchambers (i.e., > 50 % of sperm exit the microchambers in less than 10–15 seconds before washing the waste from the main channel). For devices with larger microchambers (i.e., 1000×30 μm^2^ and 500×30 μm^2^ ones), the short initial washing steps for evacuating the waste from the main channels did not have a significant effect on final progressive motile sperm concentration.

### Conclusion

We propose a new microchamber-based microfluidics platform for separation of only progressive motile sperm directly from a raw semen sample in a short period of time (~10 minutes). The loading of raw semen samples into microchambers and the subsequent retrieval of progressive motile sperm are straightforward and do not require any special skills. Once the raw semen sample, without any pre-treatment, is loaded, microchip reaches to a kinematic equilibrium state in terms of the number of sperm that enter or exit the microchambers. The equilibrium state further provides the user with flexibility and controllability of the process since there will be enough time for performing the rest of the operations. We evaluate the efficiency of our device by investigating the quality of progressive motile sperm through measuring the VSL, VAP distributions and their comparison to the original raw semen samples. We also measured and predicted the sperms’ residence time inside the microchambers as well as the whole progressive sperm collection time. In addition, we showed that the microfluidic platform not only can separate motile sperm from immotile, but also can effectively differentiate and separate progressive sperm from nonprogressive motile ones. Moreover, we optimized the concentration of the collected progressive sperm by tuning the washing flow rate above the sperm rheotaxis ability evading any further post processing. Finally, we believe that our proposed microfluidic platform can be an ideal alternative to the conventional sperm separation techniques and may contribute to the improvement of the diagnostics and ART outcome for animal and human fertility in the future.

## Materials and Methods

Frozen straws of bovine semen were thawed by immersing them in a bath of water at 37°C for 30-45 seconds immediately after being retrieved from the liquid nitrogen tank. The thawed semen were then transferred to 1.5 mL Eppendorf tubes. Then, fresh medium at 37°C was added to the sample for viability. Otherwise, the samples are not diluted. The samples were put under the microscope stage for processes less than 5 minutes. However, for the essays more than 5 minutes, a warm plate was used to ensure the viability of the sperms at 37°C and in the meantime the whole thawed sperm sample was kept on the warm plate at 37°C.

The sperm samples were treated by Tyrode’s albumin lactate pyruvate (TALP) as a fresh medium. TALP was prepared using NaCl (110 mM), KCl (2.68 mM), NaH_2_PO_4_ (0.36 mM), NaHCO_3_ (25 mM), MgCl_2_ (0.49 mM), CaCl_2_ (2.4 mM), HEPES buffer (25 mM), glucose (5.56 mM), pyruvic acid (1.0 mM), penicillin G (0.006% or 3 mg/500 mL), and bovine serum albumin (20 mg/mL) as the base solution to prepare the experimental buffer pH = 7.47.

The microfluidic chips were made of SU-8 and photolithography and poured PDMS on the resulting mold as in conventional soft lithography. Syringe pumps (New Era) were used to load the samples into the chambers at different injection rates of 0.54 mL/h and 1.24 mL/h.

The Nikon eclipse TE3000 phase contrast microscope was used to visualize the movement of the sperm in the channels. Images and video recordings were acquired at 10 frames per second with a 20× objective and a digital ANDOR Zyla 4.2 sCMOS camera. During the experiments, the microfluidic chip was kept on a heated 37 °C plate under stereo microscope for experiments more than 5 minutes. The VAP and VSL of the sperm were determined using ImageJ (version 1.52a; NIH) and resulting dot trajectories of the spermatozoa were analyzed by python 3.8. The measurement of the VAP and VSL were done on samples in a microfluidics chip with 30 μm depth to avoid the effect of channel height on the sperm velocity reported by simulations and theoretical studies before.

Fluorescent micro particles were used in producing the image of the streamlines in the main channel.

Simulations have been done in COMSOL Multiphysics 5.4a for calculation of the velocity field and shear rate. Particle tracing physics were combined with the laminar flow to track the sperm’s 2D theoretical trajectories for two different flow rates (0.7 and 1.9 μL/h) for optimizing the flow rate and investigating rheotaxis effect. The streamlines were also captured by 10 μL/h flow rate in a layout corresponding to our geometry.

## Acknowledgments

We would like to thank Farhad Javi for setting up the fluorescent particles in visualizing the streamlines in the main channels. Manufacturing process of the microfluidic platform was performed at the Cornell Nanoscale Facility (CNF), a member of National Nanotechnology Coordinated Infrastructure (NNCI), which is supported by National Science Foundation Grant ECCS-1542081.

## Conflict of interests

Authors confirm no conflict of interest.

## Supplementary Information

### 1. Optimization of the loaded volume

The basic geometry of the chip is shown in the above picture and the area confined in the red rectangle is the repetitive building block. We suppose that the confined shaded area over the area of the red rectangle should be optimized in order to maximize the volume of the loaded sample. The radius of the microchambers are assumed as R and the minimum width of the unshaded area is assumed to be *s_x_* in the x direction and *s_y_* in y direction. This width is crucial for the purpose of maintaining the intact bonding of the PDMS to glass while pressure is exerted at the inlet in the loading process. The restricted shaded area, *A_sh_*, is roughly composed of two semi-circles and two rectangles.

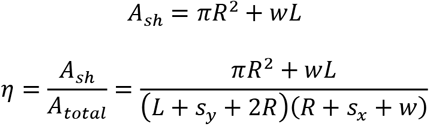

The optimization here has two restrictions which indicates the edges of the rectangle are given and have value of *χ*_0_ and *χ*_1_ for horizontal and vertical edges, respectively.

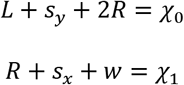

Differentiating the η and restrictions gives us the condition under which the η becomes maximum.

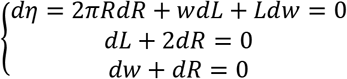

rearrangement of the equation above narrows it down to the optimal value of the microchambers’ radius.

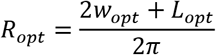

The subscript opt indicates the optimal value of the parameters here. Now by substituting this in the set of the equation of the *χ*_0_ and *χ*_1_:

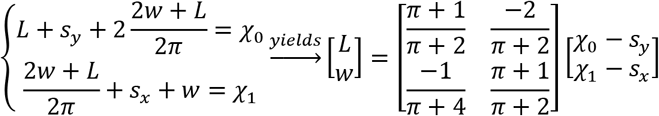

And therefore, the value of the optimum R is available with respect to χ_0,1_.

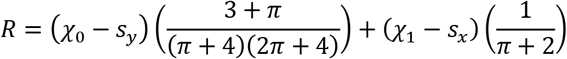

Another important restriction is that the distance between the center of the microchambers (D) should be more than the diameter of the microchambers plus a threshold for bonding considerations.

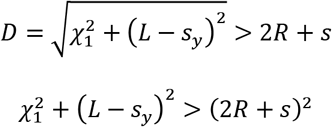

Where s is minimum distance of the microchambers’ wall with each other. Hence after designing the *χ*_0,1_, the above inequality must be checked as well.

These considerations do not take care of all other conditions important for separation. For example, one should be aware that *w* cannot be wider than tens of micrometers, or else the flow would change the stagnant condition of the microchambers during washing step. The geometries that we finally fabricated for testing the device are 6 different designs in the Table below.

### 2. SEM images of microchambers

The rough surfaces of the fabricated device may cause as random noise in the motion of sperm while swimming along the boundaries.

### 3. Theoretical Model of The Loading Time

Loading of the microchamber can be done though two different processes. First is to use constant pressure at the inlet (simply by exerting pressure on syringe plunge) and secondly, employing constant mass flow rate at the inlet. However, the main concept of the loading process remains the same. When fluid enters the microchamber the surface of the sample fluid and air trapped in the microchamber produce an angle with the walls of the microchamber which we assume to be constant and equal to the static contact angle. It holds true for a perfectly smooth wall, though due to the manufacturing randomness, the walls of the microchambers are rough. One of such a roughness and randomness of manufacturing is shown on Fig. S2 with white flash. Therefore, it is one of the model’s assumptions which result in errors against experimental data.

For both constant pressure and flow rate we presume that the main and connecting channels are filled with fluid and the solution starts as the fluid enters microchambers. Also, we assume that all microchambers are filled simultaneously with the same conditions; the fact that is roughly acceptable.

#### Constant Pressure

The pressure here means the pressure at the inlet and it differs from the air pressure trapped inside the microchamber by the amount of Laplace pressure. In this section p is the air pressure. Having said that one can write the correlation between inlet pressure and air pressure as it follows.

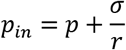

For calculation of the loaded volume (V) over time we need to figure out how θ corresponds to the V.

In triangle OO’C:

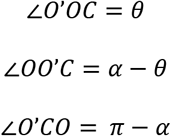

The sin law gives us:

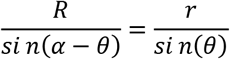

Then it can be arranged as:

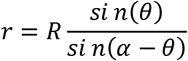

The volume of air based on the angle of θ is then calculated as

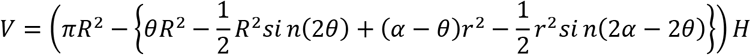

In which H is the depth of the channel and we assumed a circular common interface of air and sample fluid since H is small compared to the radius of the microchamber.

Mass conservation indicates that the rate of the change of the mass of the air in the microchamber is equal to the amount of the air running form the porous media. We then assumed that the velocity of the air expelling from the boundary of the microchamber is a function of the pressure difference of the air inside and the fresh air outside of the Polydimethylsiloxane (PDMS) device due to the Darcy’s law. The combination of the abovementioned leads to:

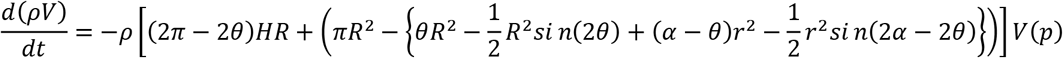

Substituting V in the previous equation and assuming a constant pressure in the air inside microchamber gives:

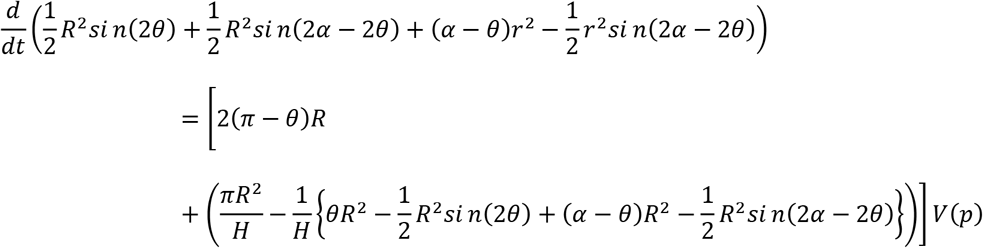

And thereafter we can get the differential of θ and r with respect to time.

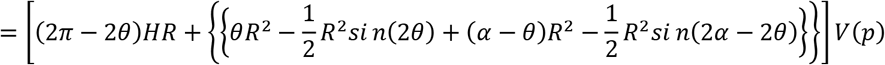

The r is the first derivative of the radius r with respect to time which is calculated as below.

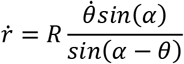

The abovementioned equation is a separable first order ordinary differential equation with initial condition of *θ* = *θ*_0_ that is the angle of the inlet of the microchamber.

#### Constant flow rate

As in the case of constant pressure the volume of the TALP inside the microchamber is

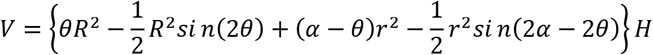

Which changes by the rate of flow inside the microchamber and r is the same as the constant pressure case.

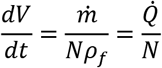

In which *ρ_f_* is the fluid density, 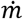 is the mass flow rate at the inlet and *N* is the total number of the microchambers.

This derivation and formula are not in the main concern of the paper and needed some extra devices to make to be validated and so we ignored about their validity. For higher depth, this formulation needs some modification regarding the interface of the sample which can no longer be assumed as a 2d curved face rather it resembles a part of a sphere surface.

#### Loading Results

Loading of the microchambers in our experiments have not been done with constant pressure nor constant mass flow rate. When we filled the main channels with the sample, we blocked the outlet and turned on the syringe pump with different flow rates. When the flow starts to fill the microchambers the pressure in the microchambers increase and then in order to keep the flow rate the pump increases the pressure at the inlet. Therefore, it is not possible to report any specific pressure at the inlet except that by increasing the flow rate the pressure increases up to a threshold in which it could be detrimental for the pump. Hence, we chose two safe flow rates of 1240 μL/h and 540 μL/hr.

### 4. Rheotaxis of the sperms in the main channel

Rheotaxis occurs when sperms come out of the microchambers and are exposed to the shear rate in the main channel resulting from the velocity gradient. The rate at which the tail of the sperm is rotated in the flow is proportional to the sinus of the angle of tail with the direction of the flow [1]. The angles and the flow direction are shown in the Fig. S4.

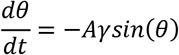

In which *γ* stands for the shear rate and A is a dimensionless empirical constant which is different for various kinds of swimmers [2]. In our case we assumed A to be 0.07.

Mathematically speaking, sperms are supposed to be dots which can swim with *v_sperm_* in the non-flow condition and therefore their trajectories is calculated by solving ordinary differential equation of lumped particles privileged by a propulsive force of their tails proportional to *v_sperm_*. The sperm’s size for bovine and human does not exceed 10 μm and hence they can be carried away by the flow from stagnant state in about 3 μs. Therefore, we assumed that the sperm velocity is simply the net resultant of fluid velocity and the velocity resulted from the motion of the flagella.

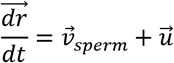

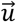 in the above equation stands for fluid velocity vector near the top surface and 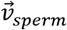 applies the propulsive velocity of sperms in its direction of movement (θ) and r shows the vector of sperm position with respect to initial position in the domain. Another assumption that will lead to equation of sperm’s trajectories is that the sperm’s average density is equal to density of the fluid otherwise the buoyancy would affect their positions.

At initial step the position of the sperm are at the point with 2.5 μm distance from the corner at each wall. *v_sperm_* are a normal distribution with mean 80 μm/s and variance 20 μm/s and *u*(*t* = 0) =80 μm/s for all of the 20 sperms at each side.

### 5. Residence time distribution model

Expansion of the boundaries of the microchamber to a congruent line of three parts with the length equal to the periphery of the microchamber and connecting channel walls. The position of each sperm on the wall is identical to its position on the line. This assumption is raised from the observations that sperms mostly remain on/near the walls (Fig. S5).

As the diameter of the microchambers increases, and more volume is loaded in the microchambers, the sperm-sperm interaction becomes more important in the calculation of the residence time, however, this effect is not considered in this model. Furthermore, as shown in the Fig. S2, some bumps and fabrication-related surface roughness, interferes with boundary navigation and speed of the sperms. Therefore, by increasing the diameter of the microchambers, the theory deviates more form experiment.

The time profile of the sperm count for different depth and different microchamber diameters are shown in the Fig. S6. The fact that the number of sperms in each microchambers differs in different trails is due to inhomogeneity of the sample. To evaluate the retrieving efficiency in the next part we counted the number of sperms being extracted from a specific row in the hope that these numbers would be consistent.

### 6. Trapping nonprogressive sperm

Nonprogressive sperm in the 1000×70 μm2 design are trapped in the microchambers and after some time (~ 15 minutes) washing crater-like hole appear in the microchambers. Fig. S7 shows these moon crater-shaped holes and how they are created by motion of nonprogressive sperm over time. The egg yolk extender particles are deviated to the outside of the circle of motion of sperm since the asymmetric motion of the tail generates a centrifugal force on the fluid that will push particles away from the center.

**Fig. S1.**
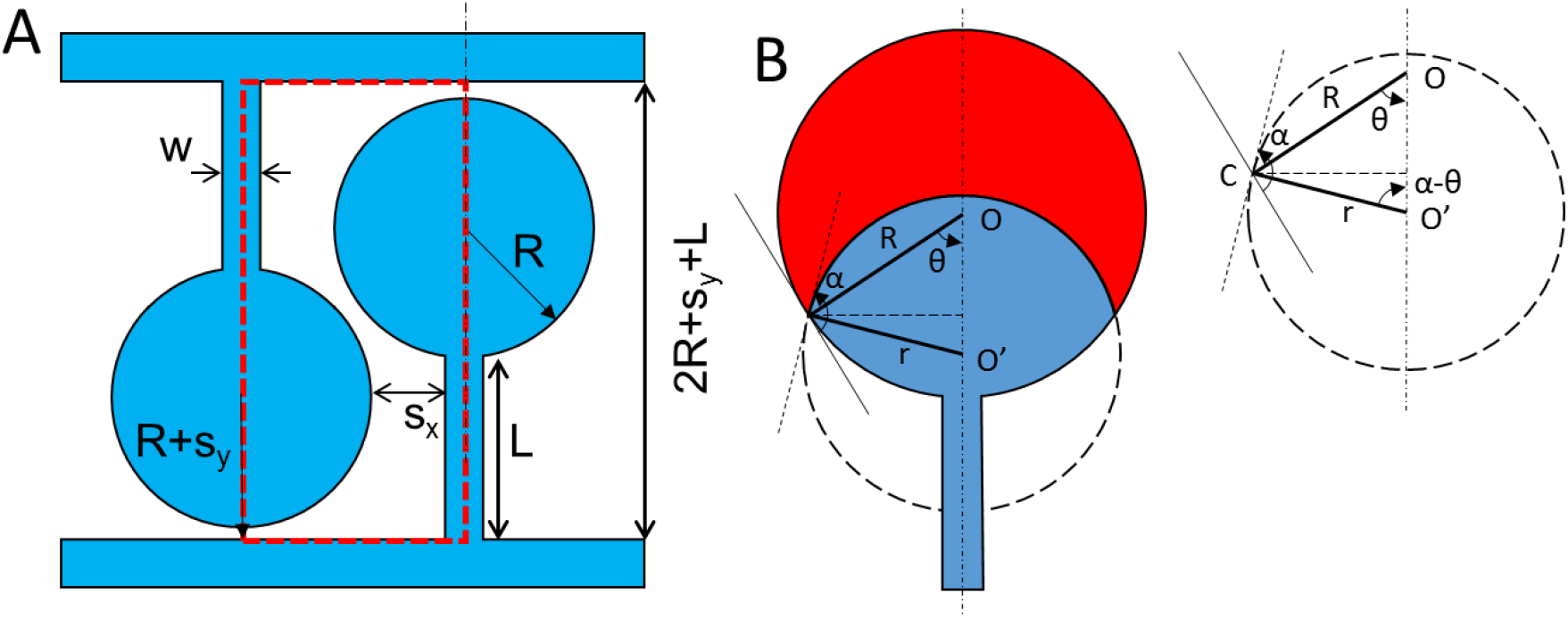
(A) The schematic of the microchannel arrays and dimensions. (B) The schematic diagram of loading process. The dimensions and parameters.

**Fig. S2.**
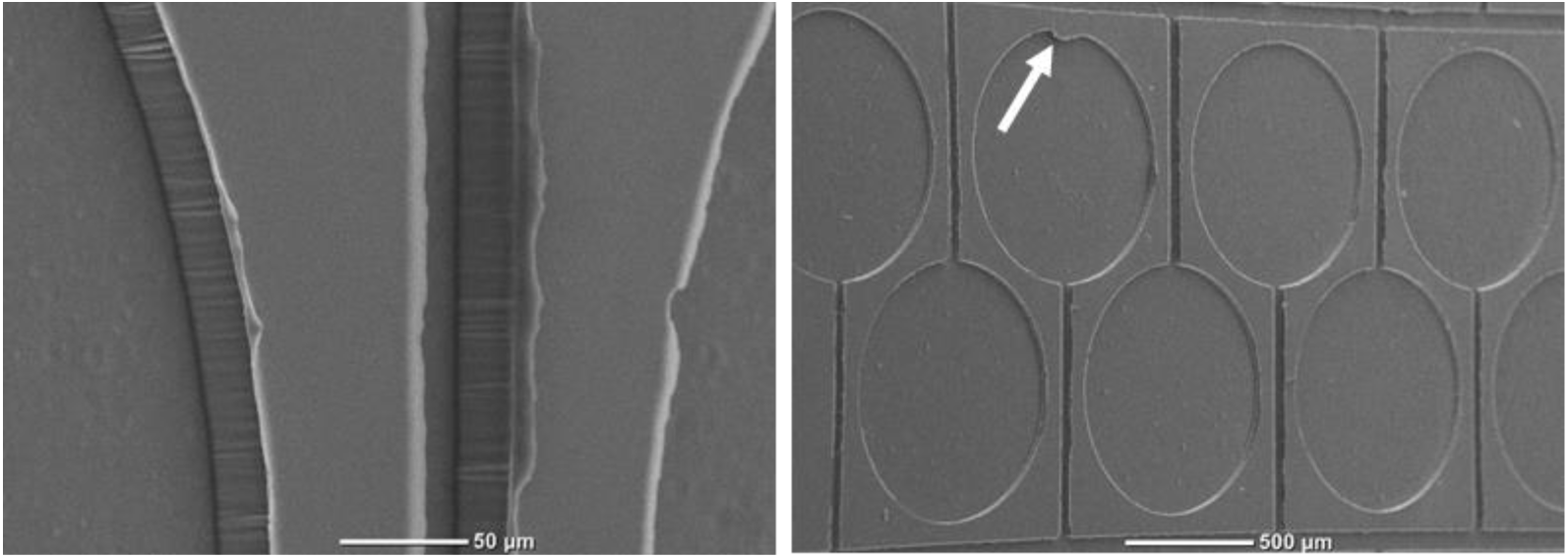
The SEM images with different magnification. The white flash shows a bump in the wall of the microchamber which cause random effects on the results of the experiments.

**Fig. S3.**
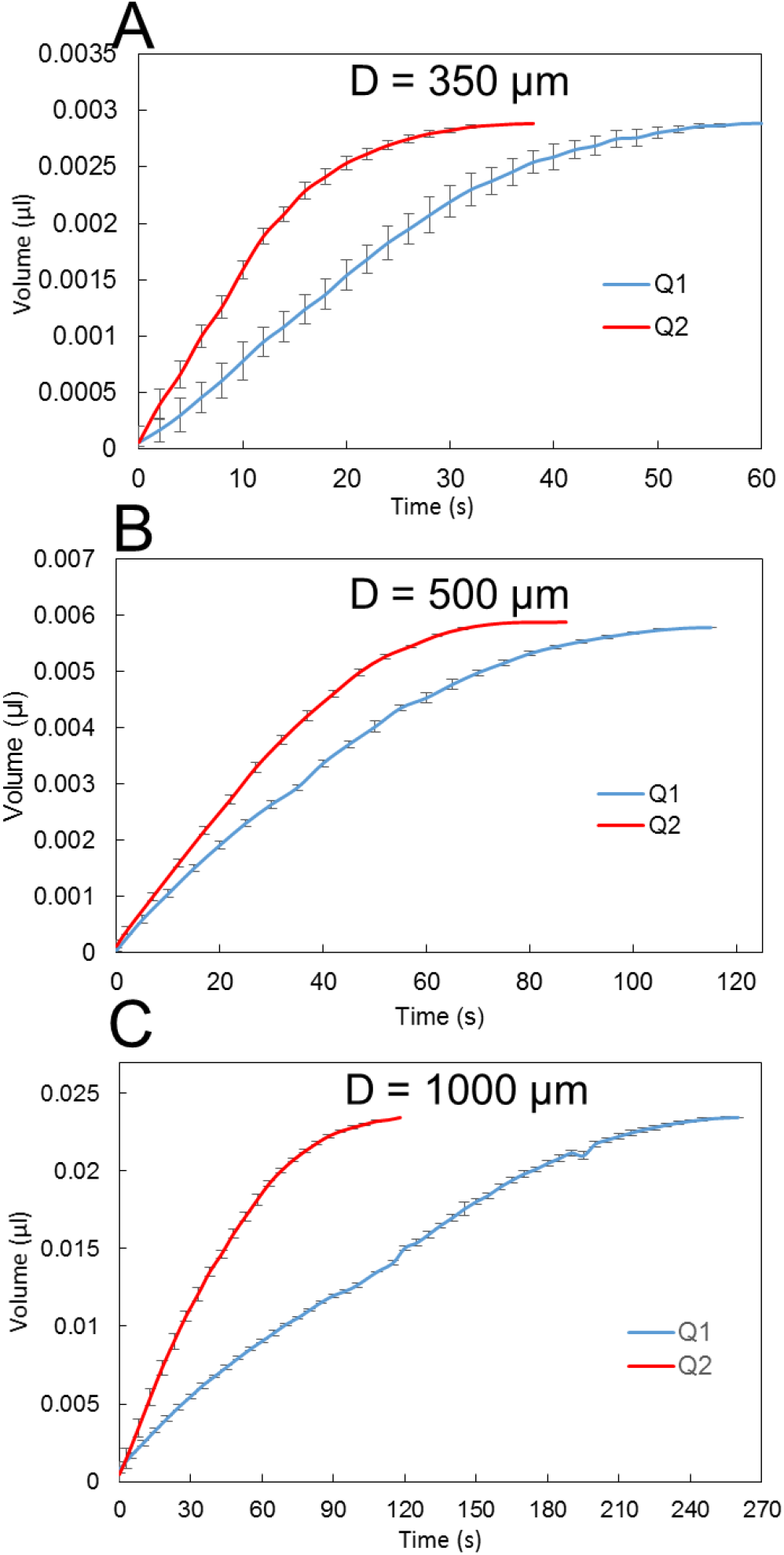
Loading volume of samples in the microchambers with different diameters for the depth of 30 μm. Q_1_=540 μL/h and Q_2_ = 1240 μL/h. (A) is for 76 microchambers, (B) is for 48 microchambers and (C) is for 36 microchambers.

**Fig. S4.**
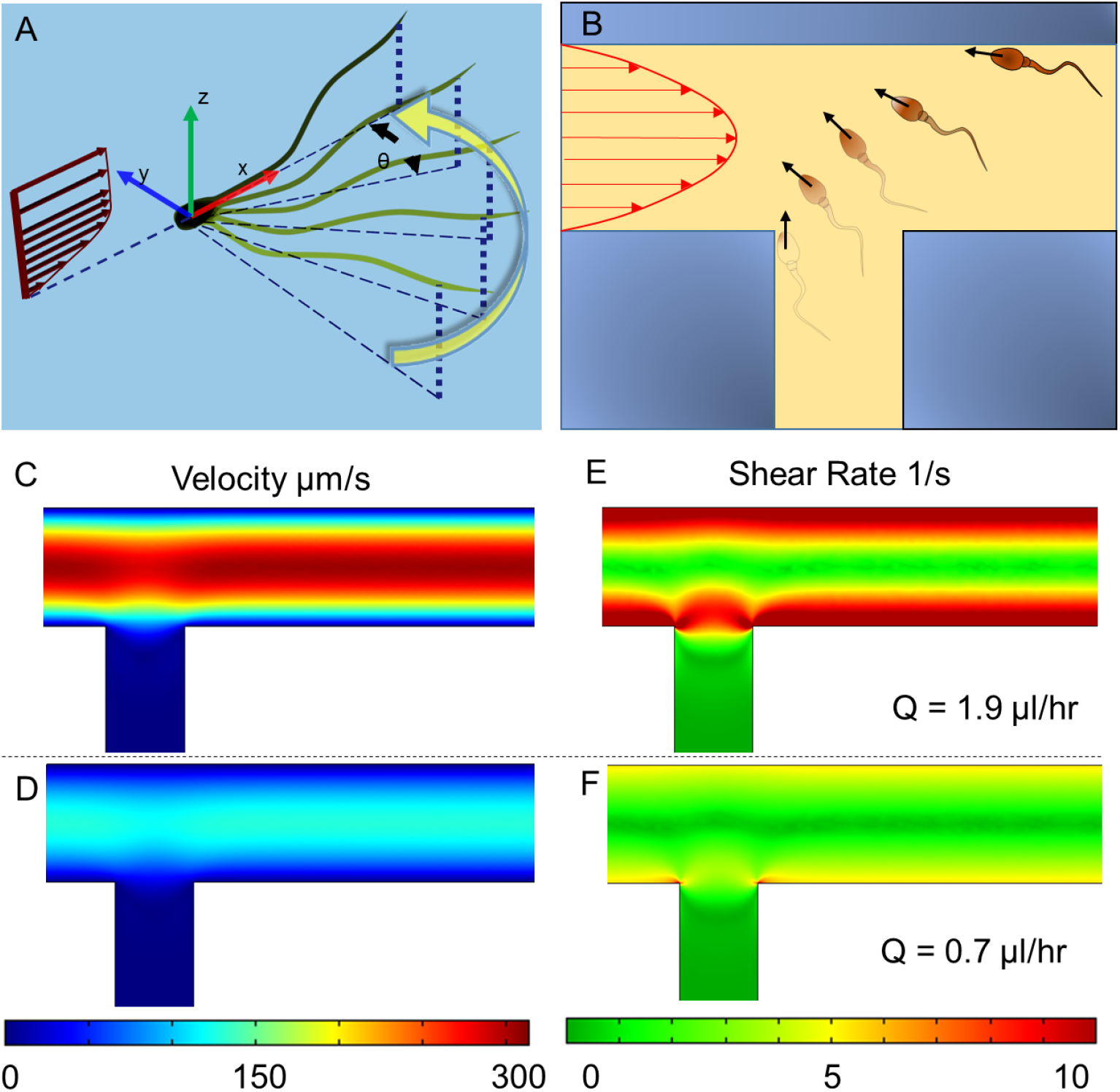
(A,B) Schematic diagrams of rheotaxis in 3d and 2d view. (C,D) the velocity magnitude and (E,F) the sear rate contours in the junction for two different flow rates in main channels with the same color map for each column.

**Fig. S5.**
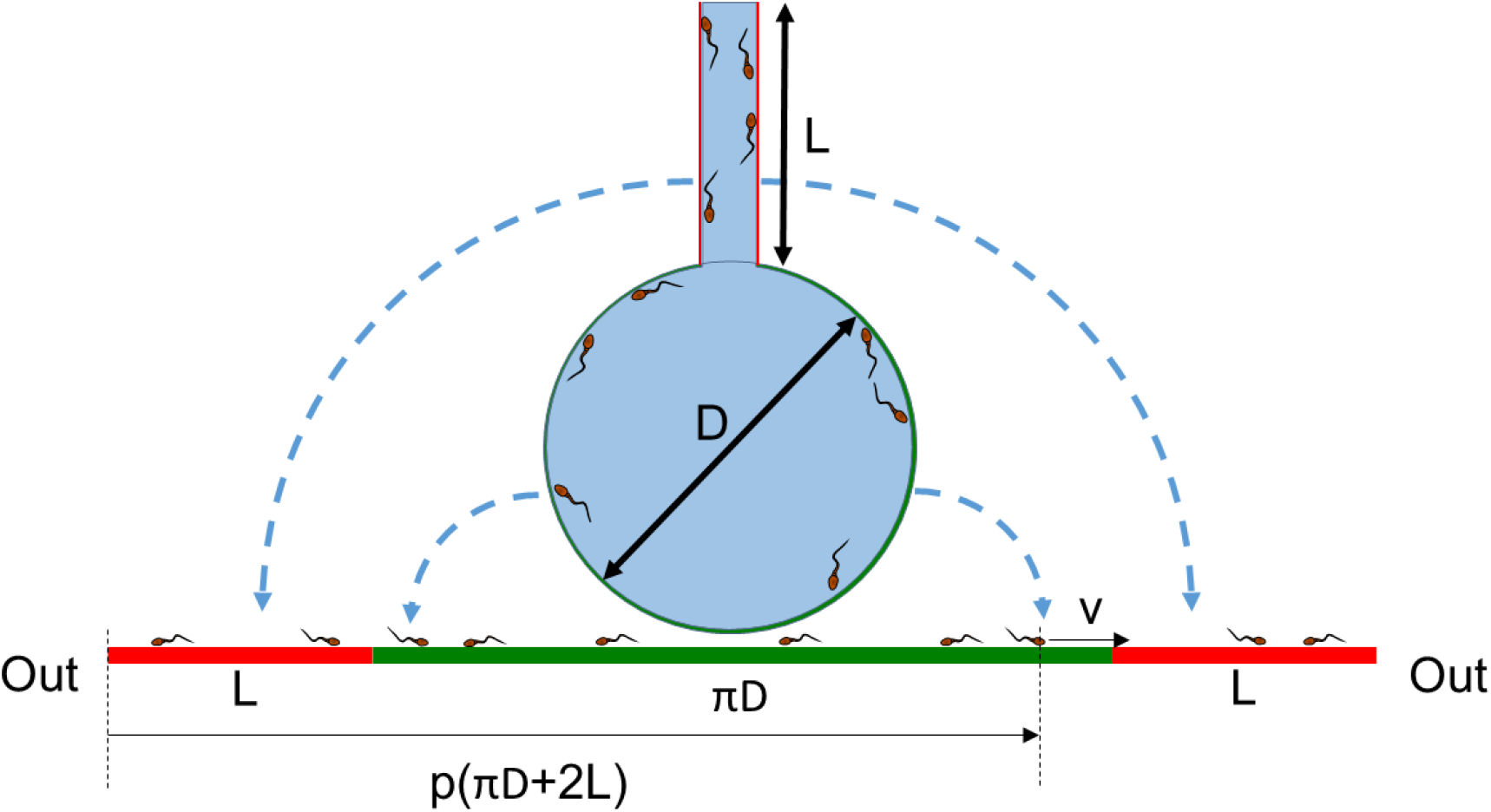
Illustration of the residence time in a congruent mapping of the microchambers boundaries on a line with same length.

**Fig. S6.**
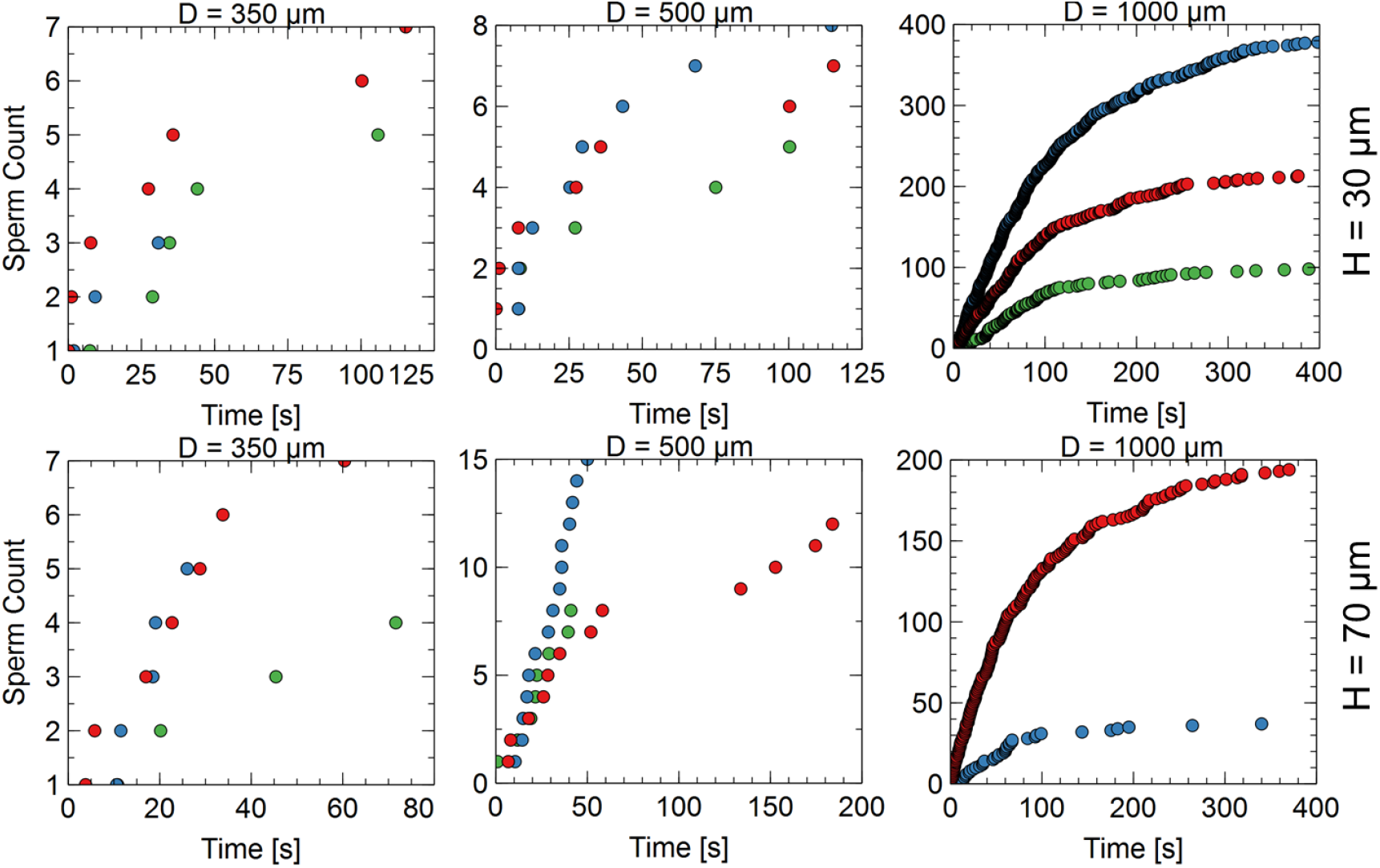
The time profile of the number of the sperms exiting the same microchamber in each diameter and depth of the chip for 3 trails. The sample concentration is 8M/mL with 13% motile sperms.

**Fig. S7.**
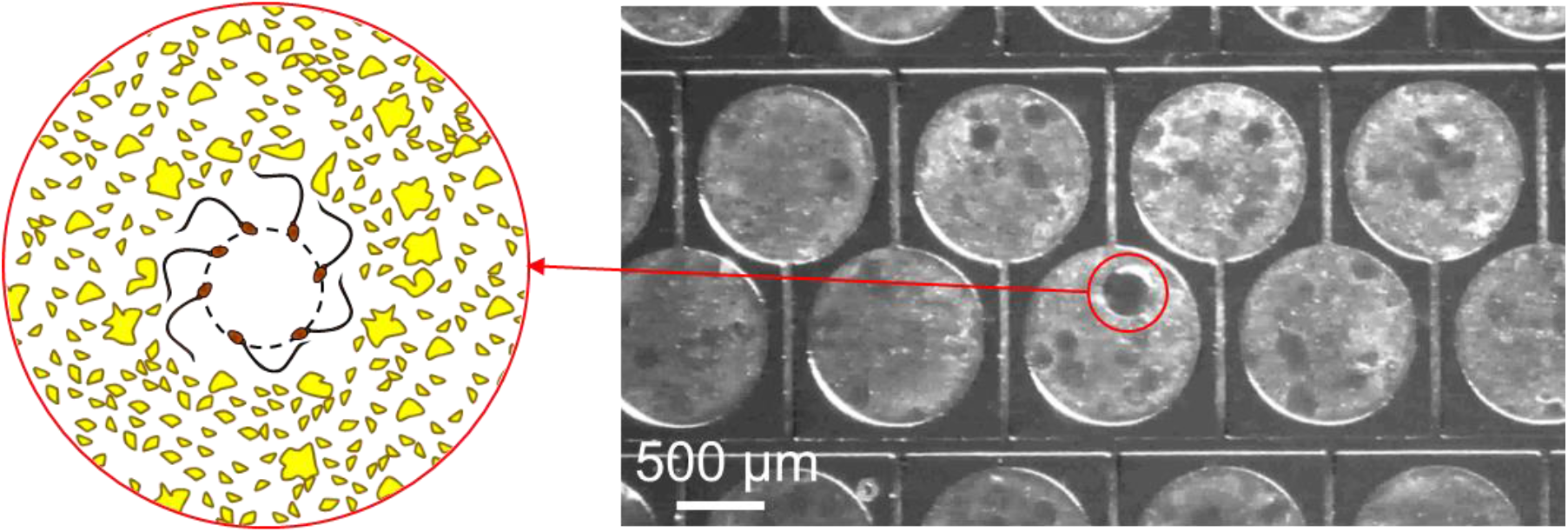
Trapping of nonprogressive spermatozoa creates moon crater-shaped hollows in the microchambers.

**Fig. S8.**
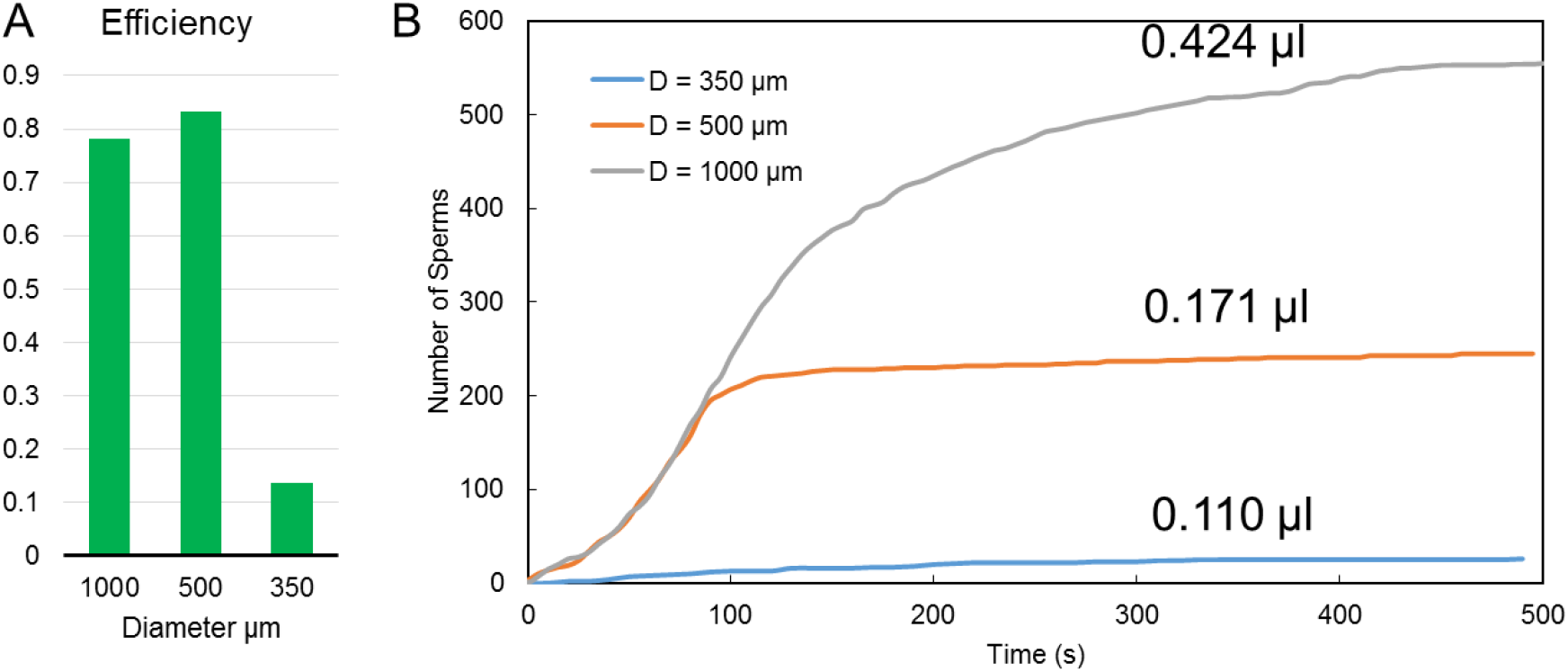
(A) Retrieving efficiency of different designs in 30 μm channel depth. (B) The diagram of the number of collected sperms from a row over time. The number of microchambers are 38, 29 and 18 for diameter of 350, 500 and 1000 μm, respectively. In this case only one side of the main channel is connected to microchambers.

**Table S1.** The dimensions of the designs used for fabrication.

| designs # | D [$\mu\text{m}$] | w [$\mu\text{m}$] | L [$\mu\text{m}$] | H [$\mu\text{m}$] | $s_x = s_y$ [$\mu\text{m}$] |
| --- | --- | --- | --- | --- | --- |
| 1 | 350 | 60 | 521.48 | 30 | 70 |
| 2 | 350 | 60 | 521.48 | 70 | 70 |
| 3 | 500 | 60 | 521.48 | 30 | 70 |
| 4 | 500 | 60 | 521.48 | 70 | 70 |
| 5 | 1000 | 60 | 978.5 | 30 | 70 |
| 6 | 1000 | 60 | 978.5 | 70 | 70 |

## Notes

### Competing Interest Statement

The authors have declared no competing interest.

